# Trisomy 21 activates the kynurenine pathway via increased dosage of interferon receptors

**DOI:** 10.1101/403642

**Authors:** Rani K. Powers, Kelly D. Sullivan, Rachel Culp-Hill, Michael P. Ludwig, Keith P. Smith, Katherine A. Waugh, Ross Minter, Kathryn D. Tuttle, Hannah C. Lewis, Angela L. Rachubinski, Ross E. Granrath, Rebecca B. Wilkerson, Darcy E. Kahn, Molishree Joshi, Angelo D’Alessandro, James C. Costello, Joaquin M. Espinosa

**Affiliations:** Linda Crnic Institute for Down Syndrome, University of Colorado Anschutz Medical Campus, Aurora, CO, USA; Computational Bioscience Program, University of Colorado Anschutz Medical Campus, Aurora, CO, USA; Department of Pharmacology, University of Colorado Anschutz Medical Campus, Aurora, CO, USA; Functional Genomics Facility, University of Colorado Anschutz Medical Campus, Aurora, CO, USA; Department of Pediatrics, University of Colorado Anschutz Medical Campus, Aurora, CO, USA; Department of Biochemistry and Molecular Genetics, University of Colorado Anschutz Medical Campus, Aurora, CO, USA; Department of Molecular, Cellular and Developmental Biology, University of Colorado Boulder, Boulder, CO, USA

## Abstract

Trisomy 21 (T21) causes Down syndrome (DS), affecting immune and neurological function by unknown mechanisms. We report here the results of a large metabolomics study showing that people with DS produce elevated levels of kynurenine and quinolinic acid, two tryptophan catabolites with potent immunosuppressive and neurotoxic properties, respectively. We found that immune cells of people with DS overexpress IDO1, the rate-limiting enzyme in the kynurenine pathway (KP) and a known interferon (IFN)-stimulated gene. Furthermore, we found a positive correlation between levels of specific inflammatory cytokines and KP dysregulation. Using metabolic flux assays, we found that IFN stimulation causes IDO1 overexpression and kynurenine overproduction in cells with T21, dependent on overexpression of IFN receptors encoded on chromosome 21. Finally, KP dysregulation is conserved in a mouse model of DS carrying triplication of the IFN receptors. Altogether, these results reveal a mechanism by which T21 could drive immunosuppression and neurotoxicity in DS.

## INTRODUCTION

Down syndrome (DS) is caused by triplication of chromosome 21 (chr21), which occurs in approximately 1 in 700 live births, representing the most common chromosomal abnormality in the human population^1^. Trisomy 21 (T21) impacts multiple organ systems during development and causes an altered disease spectrum in people with DS, significantly increasing their risk of developing Alzheimer’s disease (AD)^2^, leukemias^3^, and numerous autoimmune conditions^1,4–7^, while protecting them from solid malignancies^3,8^. However, DS-associated phenotypes are highly variable among individuals with DS, even for conserved phenotypes such as cognitive impairment and predisposition to early onset AD^9,10^. Therefore, a deeper understanding of the mechanisms driving such phenotypic variation could illuminate not only mechanisms of pathogenesis, but also opportunities for diagnostics and therapeutic strategies to serve this population. Furthermore, these mechanistic insights could also benefit larger numbers of individuals in the typical population affected by the medical conditions that are modulated, either positively or negatively, by T21.

In order to identify molecular pathways that could contribute to both the altered disease spectrum experienced by the population with DS as well as the inter-individual phenotypic variation, our group previously analyzed transcriptomes^11^ and circulating proteomes^12^ from several cohorts of individuals with DS compared to euploid individuals (D21, controls). These efforts revealed that multiple cell types from people with DS show transcriptional signatures indicative of constitutive activation of the interferon (IFN) response^11^, which could be explained by the fact that four of the six IFN receptors (IFNRs) are encoded on chr21: the two Type I IFNR subunits (*IFNAR1*, *IFNAR2*), one of the Type II IFNRs (*IFNGR2*), and *IL10RB*, which serves both as a Type III IFNR subunit and also a subunit of the receptors for IL-10, IL-22, IL-26 and IL-28^13^. Furthermore, plasma proteomics analyses demonstrated that people with DS show signs of chronic autoinflammation^12^, including elevated levels of potent inflammatory cytokines with known ties to IFN signaling (e.g. IL-6, TNF-α, MCP1). However, it remains to be defined how this obvious immune dysregulation contributes to DS-associated phenotypes, including the diverse neurological manifestations of T21. To gain further insight into this area, we investigated metabolic changes caused by T21 in the plasma of people with DS.

Here, we report the results of a large plasma metabolomics study of individuals with DS using untargeted, highly sensitive liquid chromatography mass spectrometry (LC-MS) technology. The most significantly dysregulated metabolite identified was quinolinic acid (QA), a tryptophan (TRP) catabolite with well characterized neurotoxicity, which has been repeatedly associated with diverse neurological disorders, and which is produced by activation of the kynurenine (KYN) pathway (KP)^14^. Indeed, people with DS have elevated QA, KYN and KYN/TRP ratio, all established markers of KP activation. We found that circulating immune cells of individuals with T21 show significant overexpression of indoleamine 2,3-dioxygenase 1 (IDO1), the rate-limiting enzyme in the KP and a known IFN-stimulated gene (ISG) induced by both Type I and II IFN ligands^15–17^. Using cell-based metabolic flux assays, we found that IFN-α stimulation leads to super-induction of IDO1 and hyperactivation of the KYN pathway in cells with T21, and this requires overexpression of both Type I IFNR subunits encoded on chr21. Furthermore, we found positive correlations between circulating levels of key inflammatory cytokines, including TNF-α and IFN-λ, and KP activation in people with DS. Finally, we found that a mouse model of DS carrying triplication of the IFNR gene cluster, *Dp(16)1/Yey*, shows increased levels of KYN relative to wild type littermates. Given the well-established neurotoxicity of QA, the association of KP dysregulation with wide range of neurological conditions^18–33^, as well as the immunosuppressive function of KYN^34^, our results point to KP activation as a contributing factor to neurological and immunological phenotypes in DS.

## RESULTS

### Trisomy 21 induces the kynurenine pathway

In order to identify metabolic pathways that are consistently dysregulated by T21, we collected plasma samples from two fully independent cohorts: the Translational Nexus Clinical Data Registry and Biobank (referred hereto as Cohort 1), and the Crnic Institute’s Human Trisome Project (www.trisome.org, NCT02864108, referred hereto as Cohort 2). Cohort 1 included 49 individuals with T21 and 49 euploid (D21) individuals, while Cohort 2 consisted of 26 T21 and 41 D21. The breakdown of participants according to age and sex are reported in **Supplementary Data 1**. While Cohort 1 was balanced for karyotype but imbalanced for age, Cohort 2 was more age-balanced, with a bias towards females in both groups. When the two cohorts were combined, age, sex, and karyotype are more evenly distributed, which prompted us to analyze the cohorts both individually and in combination. It should be noted that our protocol excluded participants with recent or active infections. To determine metabolite abundance, we performed untargeted, LC-MS metabolomics on plasma samples from each cohort. The combined analysis workflow is outlined in **Supplementary Fig. 1a** with details available in the **Methods**. This approach led to the identification of 91 KEGG-annotated metabolites detected in both cohorts. To identify the metabolites that were consistently differentially abundant between people with and without DS, we applied a robust linear model-fitting approach that accounted for age and sex^35^ to data from each of the cohorts (**Supplementary Data 2**). We observed both differences and overlap in the differential metabolite profiles when analyzing Cohorts 1 and 2 individually. However, due to the relative quantification by untargeted LC-MS, we observed an overall difference in the distributions of metabolite abundances between cohorts (**Supplementary Fig. 1b**). Therefore, to mitigate imbalances within the separate cohorts and to increase statistical power, we completed a combined cohort analysis by modeling age and sex covariates, along with fitting each cohort as a batch effect. Using this linear model fit, the distributions of metabolite abundance overlap and are thus comparable (**Supplementary Fig. 1c**).

**Figure 1.**
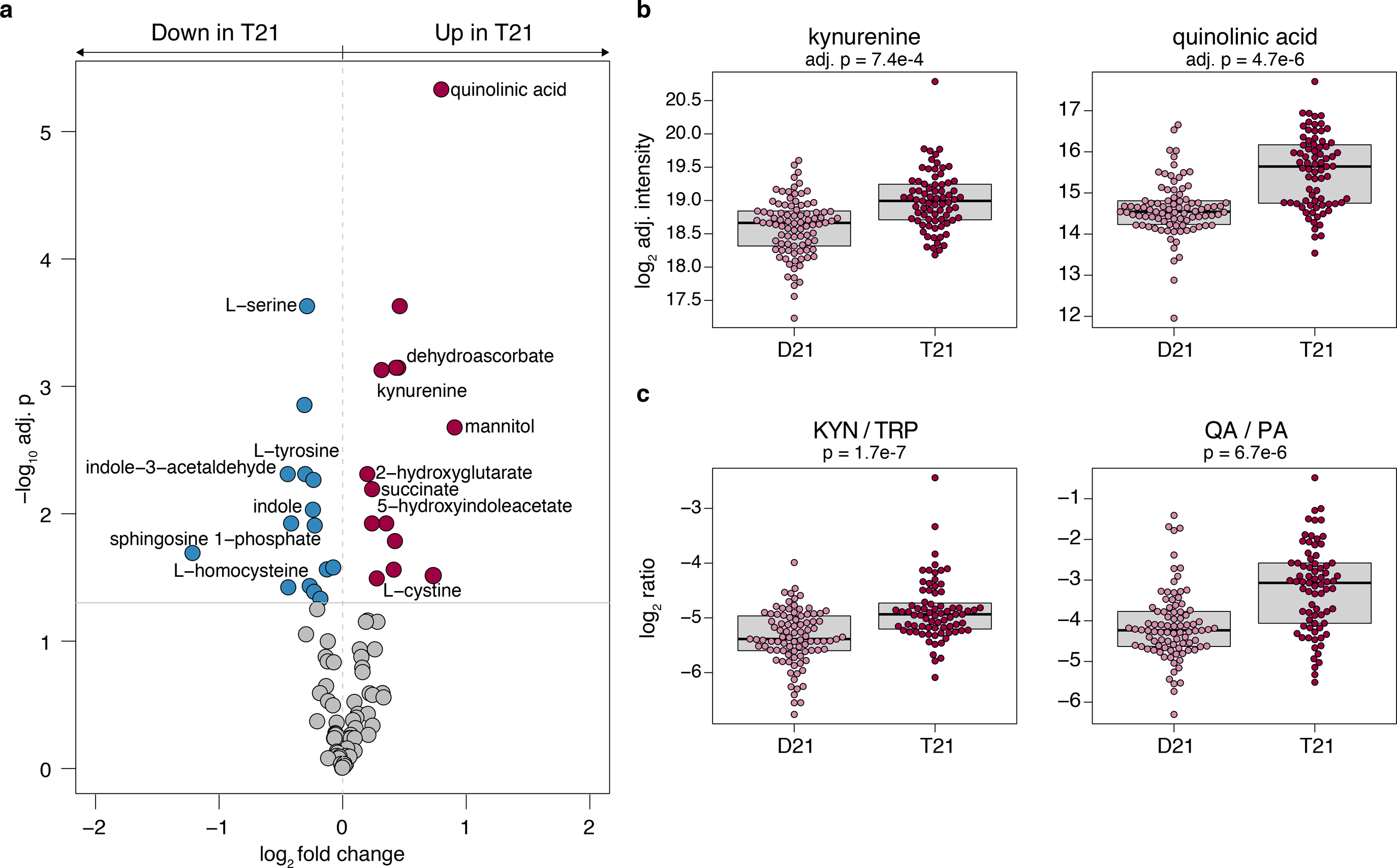
Trisomy 21 induces the kynurenine pathway. **a)** Differentially abundant metabolites in plasma samples from individuals with T21 were identified using a linear model adjusting for age, sex, and cohort. Metabolites that increased in the T21 group are shown in red and metabolites that decreased in the T21 group are blue. The horizontal grey line represents an adjusted p-value of 0.05. **b)** Boxplots showing log_2_ adjusted intensities for significantly differentially abundant metabolites. p-values were calculated using the linear model from **a** and the FDR method for multiple testing correction. **c)** Boxplots showing log_2_ adjusted ratios as indicated. p-values were calculated using an unmoderated t-test on the linear model in **a**.

When the combined total of 165 samples in the two cohorts were analyzed for the 91 metabolites using this linear model fit accounting for age, sex, and cohort, we identified 29 metabolites that were significantly differentially abundant according to a false discovery rate (FDR)-adjusted p-value on a Student’s t-test of 0.05^36^ (**Fig. 1a**, **Supplementary Data 2**).

Consistent with earlier reports^37,38^, L-serine and L-tyrosine were found to be depleted in people with DS (**Fig. 1a**, **Supplementary Fig. 1d**). Depletion of L-homocysteine (**Fig. 1a**, **Supplementary Fig. 1d**), which has also been previously observed in people with DS^39^, could be attributed to the localization of the cystathionine-beta-synthase gene (*CBS*) on chr21. Among the upregulated metabolites, we observed succinate and 2-hydroxyglutarate, two metabolites related to the TCA cycle (**Fig. 1a**, **Supplementary Fig. 1d**). Elevated succinate levels have been previously reported in people with DS and were interpreted as a sign of mitochondrial dysfunction^38^, but succinate upregulation is also an established marker of inflammation^40^. Other upregulated metabolites included the antioxidant molecules dehydroascorbate and mannitol (**Fig. 1a**, **Supplementary Fig. 1d**)^41,42^. The most significantly dysregulated metabolite in this analysis was QA, an intermediate in the TRP catabolic pathway that ultimately leads to synthesis of nicotinamide adenine dinucleotide (NAD^+^). In fact, of the 11 measured metabolites in the TRP pathway annotated by KEGG, we found five to be differentially abundant in people with DS: KYN, QA, and 5-hydroxyindoleacetate are upregulated, while indole and indole-3-acetaldehyde are downregulated (**Fig. 1a-c**, **Supplementary Fig. 2a**).

**Figure 2.**
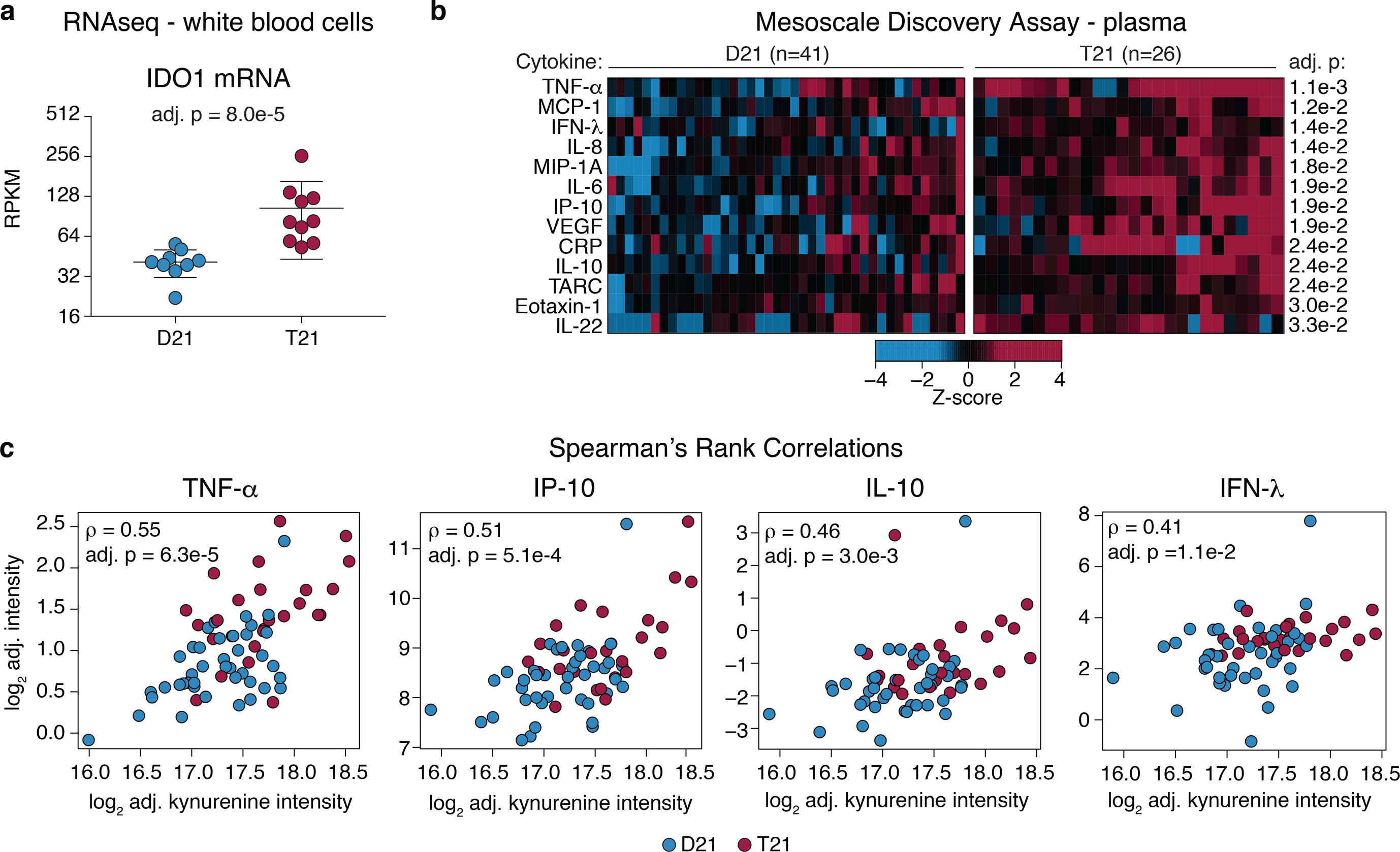
Elevated KYN levels are associated IDO1 overexpression and circulating inflammatory markers. **a)** Scatter plot showing mRNA expression of *IDO1* in white blood cells from controls (D21) and individuals with T21. Statistical significance was calculated using DESeq2. mRNA expression values are displayed in reads per kilobase per million (RPKM). **b)** Heat map showing significantly different cytokines identified in Mesoscale Discovery Assay in Cohort 2. Data from both the D21 and T21 samples were z-scored using the mean and standard deviation from the D21 group. Significant differences were assessed using the Kolmogorov-Smirnov test and an FDR-adjusted p-value threshold of 0.05. **c)** Scatter plots comparing kynurenine levels to those of the four most significantly correlated cytokines, TNF-α, IP-10, IL-10, and IFN-λ. Spearman’s rho (*ρ*) and FDR-corrected p-values are indicated for each pair.

In vertebrate cells, there are three major catabolic pathways for TRP, leading to the production of serotonin, NAD^+^, or indole (**Supplementary Fig. 2a**). Conversion of TRP to KYN represents a major pathway for catabolism of ingested TRP, eventually leading to NAD^+^ production, and accounting for removal of up to 99% of TRP not used in protein synthesis^43^. Remarkably, KYN itself, and the neurotoxic downstream product QA, are significantly elevated in people with DS (**Fig. 1a,b**). In contrast, picolinic acid (PA), a neuroprotective derivative of KYN, is not different between the two groups (**Supplementary Fig. 2a**). Accordingly, the KYN/TRP and QA/PA ratios are both significantly higher in people with DS (**Fig. 1d**).

Altogether, these results indicate that a major metabolic impact of T21 is dysregulation of TRP catabolism toward increased activity in the KP. Importantly, activation of the KP has been implicated in myriad neurological and neurodegenerative disorders, and elevation in the KYN/TRP and QA/PA ratios have been linked to the development of neurological conditions such as Alzheimer’s disease^21–24^, Huntington’s disease^25,26^, AIDS/HIV-associated neurocognitive disorder^27–29^, Parkinson’s disease^44^, amyotrophic lateral sclerosis^30,31^, and multiple sclerosis^32,33^. Additionally, KYN has potent immunosuppressive functions^34^. Therefore, we focused our efforts on elucidating the mechanisms driving activation of the KP in DS.

### Activation of the KP is associated with IDO1 overexpression and elevation of specific inflammatory cytokines

In order to investigate potential mechanisms driving the elevated levels of KYN and QA in people with DS, we performed transcriptome analysis of circulating white blood cells (WBCs) in a third cohort of 19 adult individuals, 10 of them with T21 (see **Supplementary Data 1** for information on Cohort 3). We interrogated this dataset to assess possible alterations in the expression levels of various enzymes catalyzing different reactions in the KP (see **Supplementary Fig. 2a,b**). This exercise revealed a significant increase in the expression level of IDO1, the enzyme that catalyzes the rate-limiting step in the KP, in people with DS (**Fig. 2a, Supplementary Data 3**). The only other significant change observed was a decrease in kynurenine 3-monooxygenase (KMO), the enzyme that converts KYN into 3-hydroxykynurenine (**Supplementary Fig. 2a-c)**. Given that *IDO1* is a well-characterized ISG, known to be stimulated by both Type I and II IFN signaling^15–17^, this systemic IDO1 overexpression could be explained by our earlier finding that different immune cell types from people with DS show consistent hyperactivation of the IFN response^11^. Thus, these results indicate that the observed activation of the KYN pathway in people with DS could be explained simply by higher levels of IFN signaling and IDO1 overexpression.

We recently reported the results of a large plasma proteomics study of people with DS^12^, which revealed that T21 causes changes in the circulating proteome indicative of chronic autoinflammation, including elevated levels of many cytokines acting downstream of IFN signaling. Therefore, we tested whether there was a correlation between levels of inflammatory cytokines and KP by using a Mesoscale Discovery (MSD) assay to measure a panel of 55 cytokines in the plasma samples from the 67 participants in Cohort 2. First, this analysis confirmed the previously reported upregulation of several cytokines in people with DS, including TNF-α, MCP-1, IL-6, VEGF, and IL-22 (**Fig. 2b**, **Supplementary Data 4**)^12^. Furthermore, we identified several additional inflammatory markers as dysregulated in individuals with T21, including C-reactive protein (CRP) and IFN-λ, also known as IL-29 (**Fig. 2b**, **Supplementary Data 4**). To investigate the relationship between KYN and each of the cytokines measured, we calculated Spearman’s rank correlations and found ten significant correlations (**Supplementary Data 5**). At the top of this list were three cytokines known to be activated downstream of IFN, the proinflammatory molecules TNF-α and IP-10 (IFN-γ induced protein 10, CXCL10), as well as the anti-inflammatory cytokine IL-10 (**Fig. 2c**, **Supplementary Data 5**). Furthermore, the fourth strongest correlation with KYN levels was the Type III IFN ligand IFN-λ (**Fig. 2c**, **Supplementary Data 5**), which is interesting as IFN-λ has also been shown to induce expression of IDO1^45^.

Taken together, these data indicate that increased KYN levels are associated with IDO1 overexpression and increased levels of specific inflammatory markers *in vivo*, consistent with constitutive IFN activation and immune dysregulation in DS.

### Trisomy 21 sensitizes cells to super-induction of the KP via increased IFNR gene dosage

Given the wealth of potential mechanisms that could alter TRP catabolism in individuals with DS, including differences in medical histories, existing co-morbidities, dietary regimes, and medication intake, we asked whether KP dysregulation could be observed at the cellular level. Toward this end, we employed cell-based metabolic flux experiments using heavy isotope-labeled (^13^C^15^N) TRP on a panel of age- and sex-matched skin fibroblasts derived from individuals with and without T21. Given the widely-established fact that the KP is activated by Type I and II IFN signaling^15–17^, we chose to perform these experiments before and after IFN-α stimulation. Matched western blot analysis showed that T21 fibroblasts display much stronger induction of IDO1 protein expression relative to D21 cells (**Fig. 3a-b**). Remarkably, IFN-α stimulation produced a significant, time-dependent depletion of the isotopic TRP in T21 cells, but not supernatant (**Fig. 3c**, **Supplementary Fig. 3a, Supplementary Data 6**), concurrent with a significant increase in KYN levels in the supernatant, but not in cells (**Fig. 3d**, **Supplementary Fig. 3b**). Accordingly, the KYN/TRP ratios were most significantly elevated upon IFN-α stimulation in T21 cell cultures, consistent with more pronounced consumption of intracellular TRP and subsequent secretion of elevated levels of KYN in cells of people with DS (**Fig. 3e**, **Supplementary Fig. 3c,d**).

**Figure 3.**
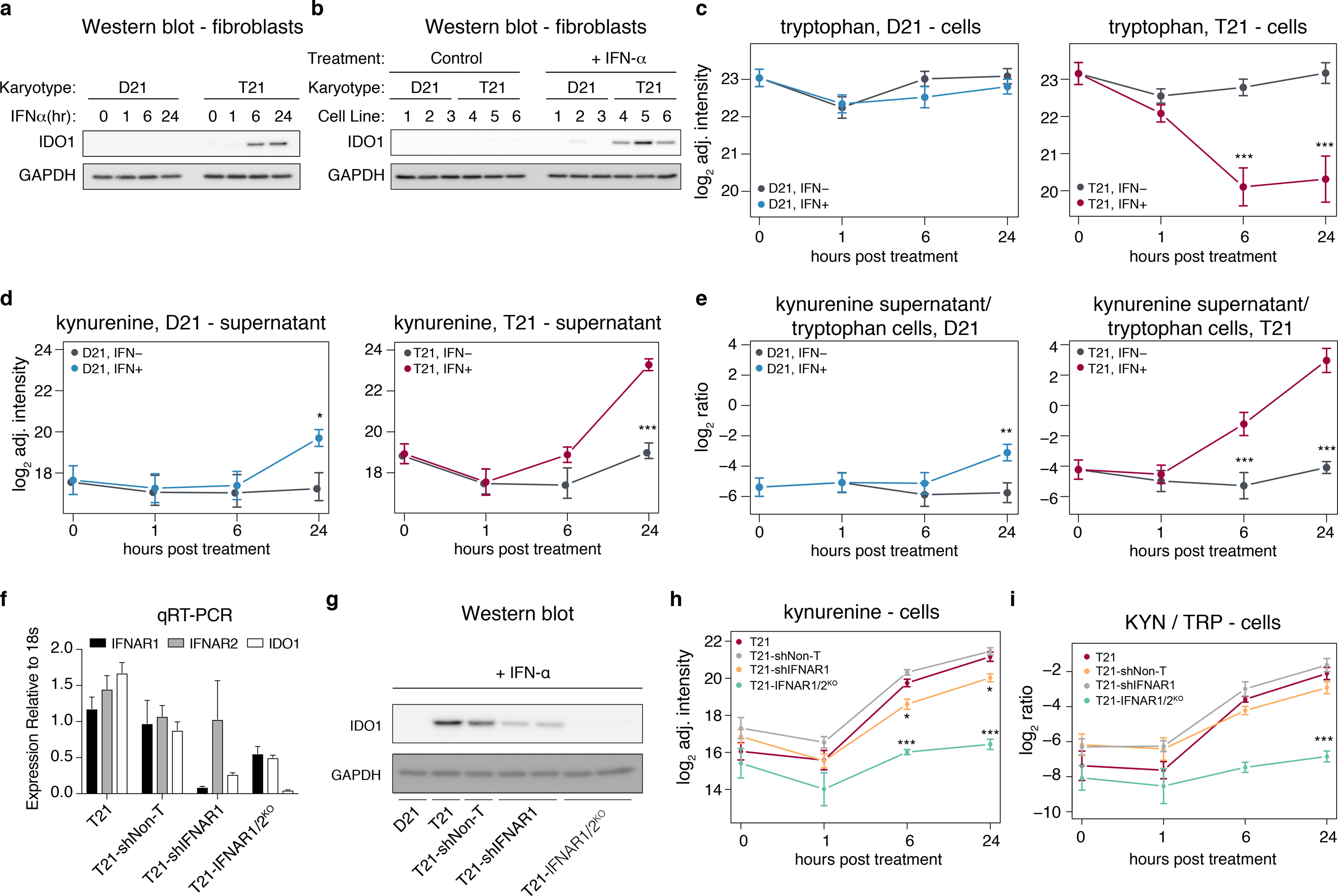
Trisomy 21 sensitizes cells to super-induction of the kynurenine pathway by IFN-α. **a)** Western blot analysis demonstrating upregulation of IDO1 over a 24 hour time course in an age- and gender-matched pair of fibroblasts with (T21) or without (D21) trisomy 21. **b)** Western blot analysis confirming IDO1 super-induction in multiple T21 fibroblasts after 24 hours of IFN-α treatment. 1-3 are cell lines from typical people, 4-6 are cell lines from age- and gender-matched individuals with trisomy 21. **c)** Metabolic flux experiment using heavy-labeled (^13^C^15^N) tryptophan in fibroblast cell lines (n = 3 T21, n = 3 D21). Levels of heavy-labeled tryptophan in D21 (left) and T21 (right) fibroblast cell lysates, with or without IFN-α treatment. **d)** Levels of heavy-labeled kynurenine in D21 (left) and T21 (right) supernatant, with or without IFN-α treatment. **e)** Ratio of heavy-labeled kynurenine in supernatant to heavy-labeled tryptophan levels in cells in D21 (left) and T21 (right) supernatant, with or without IFN-α treatment. At each timepoint, the p-value between the Ctrl and IFN+ samples was calculated using a two-tailed student’s t-test (* p < 0.05, ** p < 0.01, *** p < 0.001). **f**. Q-RT-PCR showing relative levels of *IFNAR1*, *IFNAR2*, and *IDO1*, in the indicated cell lines. T21-shNon-T indicates cell line expressing a non-targeting shRNA. T21-shIFNAR1 indicates cell line expressing a shRNA targeting IFNAR1. T21-IFNAR1/2KO indicates a cell population transduced with guide RNAs targeting the first exons of IFNAR1 and IFNAR2. mRNA expression is expressed relative to 18s ribosomal RNA. **g**. Western blot showing effects of modulating *IFNAR* levels as in **f** on induction of IDO1 after six hours of IFN-α stimulation. **h**. Levels of heavy-labeled kynurenine in the indicated cell lines during a 24 hour time course of IFN-α treatment. **i**. Ratio of heavy-labeled kynurenine to heavy-labeled tryptophan levels in the indicated cell lines during a 24 hour timecourse of IFN-α treatment. At each timepoint, the p-value between the parental T21 cell line and each T21 cell line with modified *IFNAR* levels was calculated using a two-tailed student’s t-test (* p < 0.05, ** p < 0.01, *** p < 0.001).

Next, we tested whether the observed super-induction of IDO1 and the KP by IFN in T21 cells is due to the extra copy of the Type I IFNRs encoded on chr21 (*IFNAR1*, *IFNAR2*), using both shRNA-mediated knockdown and CRISPR-based knock-out approaches. First, we stably knocked down *IFNAR1* expression in one of the T21 cell lines by >90% with a specific shRNA that did not affect *IFNAR2* expression (T21-shIFNAR1, **Fig. 3f**). Knocking down *IFNAR1* resulted in a reproducible decrease in IFN-α-induced IDO1 expression at both the mRNA and protein levels (**Fig. 3f,g**). Independently, we also co-transduced two independent guide RNAs (gRNAs) targeting the first exons of *IFNAR1* and *IFNAR2*, which created a population of T21 fibroblasts expressing ~50% of both the *IFNAR1* and *IFNAR2* mRNAs relative to the parental cell line (T21-*IFNAR1/2*^KO^, **Fig. 3f**). Notably, such reduction in the expression of both Type I IFNRs completely abolished overexpression of IDO1 mRNA and protein in response to IFN-α stimulation, bringing it down to levels comparable to those observed in the D21 cell line (**Fig. 3f,g**). We then repeated the metabolic flux experiment with isotope-labeled TRP on this panel of T21 cell lines with reduced IFNR expression during a 24-hour time course of IFN-α treatment. Remarkably, the T21-*IFNAR1/2*^KO^ cell line showed significantly reduced levels of KYN in both cells and supernatants, as well as significantly reduced KYN/TRP ratios (**Fig. 3h-i**, **Supplementary Fig. 3e-f**). Furthermore, the T21-shIFNAR1 cell line showing intermediate IDO1 expression displayed a partial rescue of elevated KYN levels in cells, but not supernatants (**Fig. 3h**, **Supplementary Fig. 3f**).

Altogether, these results indicate that IDO1 induction by IFN-α is highly sensitivity to *IFNR* dosage, providing a mechanism by which T21 could sensitize cells to IFN-dependent super-activation of the KP. These results also demonstrate that KP dysregulation is observed at the cellular level, indicating that the elevated levels of KYN and QA in the circulation of people with DS is likely driven by the trisomy, rather than other variables.

### Tryptophan catabolism is selectively disrupted in the *Dp(16)1/Yey* mouse model of DS

To test the impact of various groups of genes encoded on human chr21 on TRP metabolism, we employed three cytogenetically distinct mouse strains carrying segmental duplications syntenic to human chr21^46^: *Dp(10)1/Yey* (Dp10), *Dp(16)1/Yey* (Dp16), and *Dp(17)1/Yey* (Dp17), which have triplication of ~35 genes from mouse chr10, ~120 genes from chr16, and ~20 genes from chr17, respectively, and compared them to wild type C5Bl/6 mice (WT) (**Fig. 4a)**. We prepared plasma samples from age- and sex-matched cohorts comprised of these strains and measured KYN using LC-MS. Interestingly, we found that the only strain with significantly increased basal plasma KYN levels relative to WT mice was the Dp16 model, which is the largest segmental duplication, and the one harboring the four IFNRs homologous to those found on human chr21: *Ifnar1*, *Ifnar2*, *Ifngr2*, and *Il10rb* (**Fig. 4a,b**, **Supplementary Data 7**). Furthermore, we and others have previously shown that not only are the IFNRs overexpressed in this model, but downstream IFN signaling is chronically activated on numerous tissues including blood and brain^11,47^. These results are consistent with a genetic basis for the observed dysregulation of the KYN pathway, and further implicate triplication of the IFNR locus as a key contributor.

**Figure 4.**
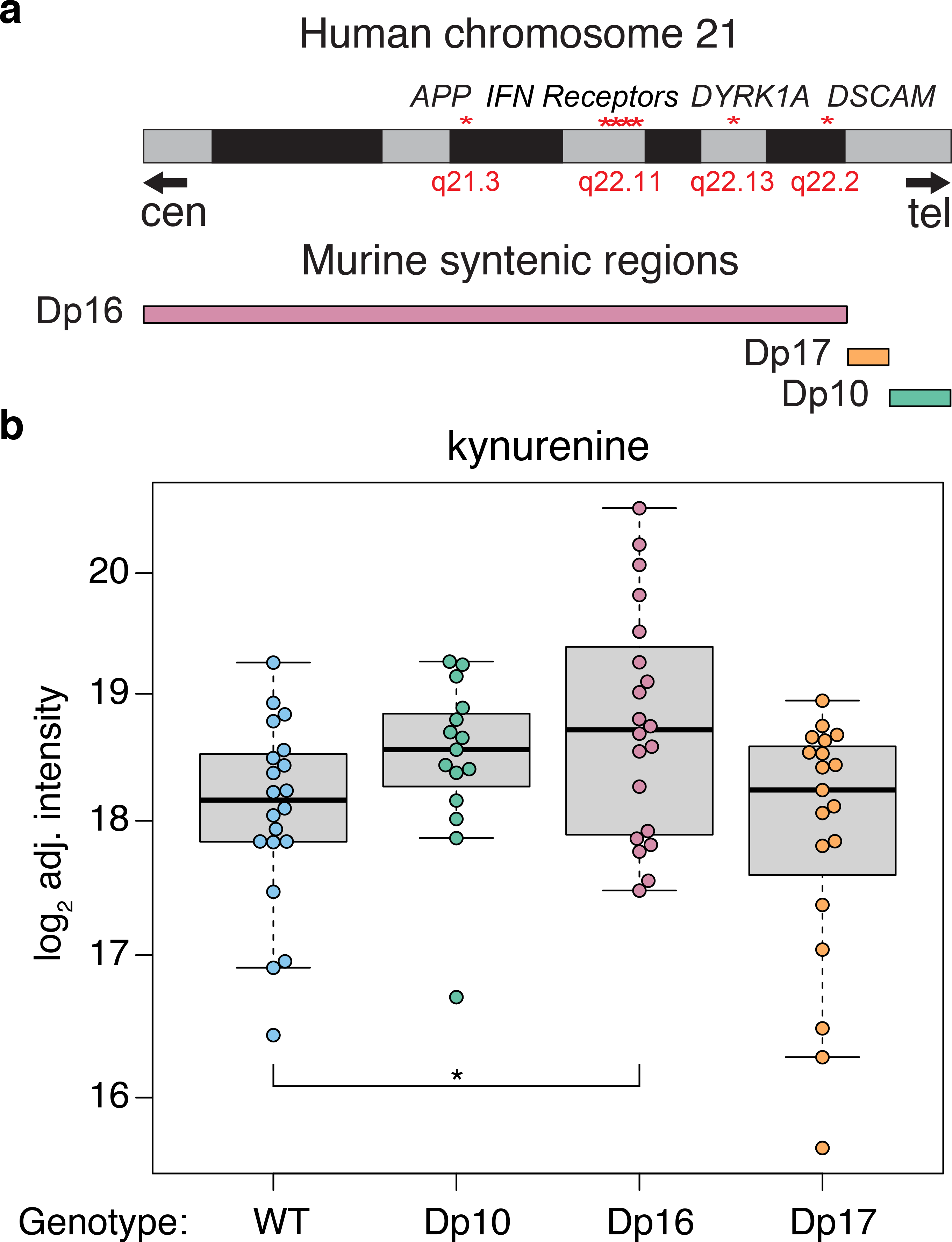
Tryptophan catabolism is disrupted in the *Dp(16)1/Yey* mouse model of DS. **a)** Schematic of mouse models tested. **b)** Box and whisker plots showing levels of plasma kynurenine in the indicated mouse model. p-values were calculated using a two-tailed student’s t-test (* p < 0.05).

## DISCUSSION

The developmental and clinical impacts of T21 in the population with DS are diverse and highly variable, with many co-morbidities showing increased prevalence in this population relative to typical people. Despite many research efforts in this area, little is known about the molecular pathways that drive the various co-morbidities common in DS. Clearly, identification of these pathways could facilitate the design of diagnostic and therapeutic strategies to serve this at-risk population via precision medicine approaches. An obvious impact of these approaches would be in the management of diverse neurological conditions showing increased prevalence in this population, including, but not restricted to, early onset AD^2^, epilepsy^48^, depression^49^, and autism^50^. Another key area of interest is in the management of immune-related disorders. On one hand, people with DS show increased incidence of autoimmune disorders such as autoimmune hypothyroidism^51^, celiac disease^7^, and myriad autoimmune skin conditions^52^, indicating a hyperactive immune system. Conversely, people with DS show increased risk to specific bacterial infections, with bacterial pneumonia being a leading cause of death^53^. Our results point to dysregulation of the KP as a potential contributing factor to the development of some neurological and immunological phenotypes in people with DS, with obvious diagnostics and therapeutic implications as described below.

People with DS constitute the largest population with a strong predisposition to AD, which is attributed to the fact that the *APP* gene is located on chr21^2^. By age 40, virtually all individuals with DS present the brain pathology of AD; however, the age of dementia diagnoses is highly variable, with some individuals being diagnosed in their 30’s, while others remain dementia-free in their 70’s^10^. Therefore, the population with T21 provides a unique opportunity to identify mechanisms that modulate the impact of amyloidosis to either accelerate or delay AD development. Interestingly, the KP has been consistently implicated in AD progression. Multiple studies have reported elevated levels of metabolites in the KP in people with AD, and, in many of these studies, KP dysregulation has been positively correlated with AD progression (reviewed in^54^). Furthermore, IDO1 inhibition reversed AD pathology in a mouse model of AD^55^. Based on these observations, it is possible to hypothesize that individuals with DS showing the strongest degree of dysregulation in the IFN→IDO1→KYN axis may also display earlier onset and faster progression of AD. Clearly, testing this hypothesis will require a significant effort, including a multi-dimensional, longitudinal analysis of IFN signaling, KP dysregulation, and various metrics of the AD pathological cascade in people with DS.

Beyond AD, it is possible that KP dysregulation is a contributor to other neuropathologies more prevalent in DS, such as epilepsy/infantile spasms/seizures, depression, and autism^48–50^. KYN and QA were shown to promote seizures in mice, rats, and frogs, and QA was found to be elevated in mouse models of epilepsy, leading to the characterization of QA as an ‘endogenous convulsant’ with a pathogenic role in seizure disorders (reviewed in^56^). The role of TRP catabolism in depression is undisputed, with a prominent role for serotonin depletion as a causative event, leading to the widespread therapeutic use of selective serotonin reuptake inhibitors (SSRIs). Importantly, activation of the KP shunts TRP away from serotonin synthesis, causing serotonin depletion, which led Lapin and others to propose the ‘kynurenine hypothesis’ of depression^57^. Furthermore, induction of the KP could not only drive serotonin deficiency, but also promote depression-associated anxiety, psychosis, and cognitive decline^58^. Although we did not observe a significant depletion of serotonin in our research participants with DS (**Supplementary Fig. 2a**), future studies could investigate whether individuals with DS diagnosed with depression show increased activation of the KP concurrent with serotonin depletion. Notably, it has been amply documented that therapeutic administration of IFN-α can induce depression, leading Wichers et al. to propose that IDO1 induction (and consequent neurotoxicity) was the principal pathophysiological mechanism^17^. Regarding autism, independent cohort studies have documented dysregulation of KYN metabolism in patients diagnosed with autism spectrum disorder^59,60^, as demonstrated by increased KYN/kynurenic acid ratios^60^ and elevated levels of QA^59^. The higher prevalence of autism in people with DS, coupled to our finding of increased levels of KYN in plasma from people with DS, support the growing notion of neuroinflammatory processes in the etiology of autism spectrum disorders.

In addition to the recognized neurological conditions discussed above, KP dysregulation could contribute to the cognitive deficits associated with DS. The acute effects of KP metabolites, QA in particular, on cognitive function are well-documented^14,61^. QA is a potent neurotoxin that acts as an excitotoxic *agonist* of NMDA receptors. Importantly, KP dysregulation in DS could explain a large body of work showing beneficial effects of memantine, an NMDA *antagonist* that protects from QA-mediated neurotoxicity, in animal models of DS^62–64^. Of note, memantine is an approved drug for the treatment of AD, which is currently being tested in a follow-up Phase II clinical trial to improve cognition in young adults with DS (NCT02304302). The initial pilot trial showed benefits on a subset of neurocognitive measures^65^, and a second independent trial, which focused on adults 40 years and older, showed no effects in any of the endpoints^66^. However, we posit here that the limited therapeutic effect of this drug is due to its action at a very downstream step in the pathway, and that inhibitors of IDO1 or IFN signaling (e.g. JAK inhibitors) should be considered.

Finally, it is important to note that the KP plays major roles in immune control. KYN is a potent immunosuppressive metabolite promoting ‘immune privilege’ in tissues such as the brain, testes, and gut, but which is also exploited by some tumor types to evade immune surveillance^34^. Hyper-activation of the KP in people with DS upon immune stimuli may affect key cell types involved in autoimmunity, such as regulatory T cells, and it may represent an immunosuppressive negative feedback mechanism downstream of exacerbated IFN signaling. Given that people with DS are highly predisposed to myriad autoimmune disorders, and that *pneumonia* lung infections are a leading cause of death in people with DS^53^, our results justify future investigations to define the role of KP dysregulation in multiple common co-morbidities in DS.

## Materials and Methods

### Human cohorts and sample collection

All human subjects in this study were consented according to Colorado Multiple Institutional Review Board (COMIRB)-approved protocols. Written informed consent was obtained from parents or guardians of participants under the age of 18, and assent was obtained from participants over the age of 7 who were cognitively able to assent. Deidentified plasma samples were obtained from the Translational Nexus Clinical Data Registry and Biobank (University of Colorado Anschutz Medical Campus, COMIRB #08-1276) for 49 individuals with DS and 49 controls without DS. Additionally, plasma samples for 26 individuals with DS and 41 controls were obtained through the Crnic’s Institute Human Trisome Project (University of Colorado Anschutz Medical Campus, COMIRB #15-2170, www.trisome.org). Plasma was collected in Vacutainer tubes (EDTA – purple capped or Lithium heparin – light green capped) and stored at −80°C. Participant medical history was collected by the respective biobanks. A third cohort of 19 individuals (10 with T21), was recruited through the Human Trisome Project for RNAseq analysis of white blood cells (WBCs).

### Metabolomic sample preparation

For plasma analyses (human and mouse plasma), a volume of 50 μl of was extracted in 450 μl of ice cold lysis buffer (methanol:acetonitrile:water 5:3:2). For cell-based experiments, 1×10^6^ cells were extracted in 1 ml of lysis buffer. In both protocols, incubation with lysis buffer was followed by vortexing for 30 minutes at 4°C. Separate extractions for hydrophobic metabolites (including oxylipids) were performed in ice cold methanol with similar ratios and workflows. Insoluble proteins were pelleted by centrifugation (10 minutes at 4°C and 10,000 g) and supernatants were collected and stored at −80°C until analysis. UHPLC-MS metabolomics analyses were performed using a Vanquish UHPLC system coupled online to a Q Exactive mass spectrometer (Thermo Fisher, Bremen, Germany). Samples were resolved over a Kinetex C18 column (2.1 × 150 mm, 1.7 μm; Phenomenex, Torrance, CA, USA) at 25°C using a three minute isocratic condition of 5% acetonitrile, 95% water, and 0.1% formic acid flowing at 250 μl/min or using a 5 min gradient at 400 μl/min from 5-95% B (A: water/0.1% formic acid; B: acetonitrile/0.1% formic acid) and a 17 min gradient ((A) water/acetonitrile (75/25, vol/vol) with 5 mmol/L ammonium acetate and (B) 2-propanol/acetonitrile/water (50/45/5, vol/vol/vol) with 5 mmol/L ammonium acetate). Compounds of interest were monitored and quantified in negative ion mode against deuterium-labeled internal standards. MS analysis and data elaboration were performed as described^67,68^. To monitor possible technical variability, technical replicates were run for both cohorts. Briefly, for the plasma samples from each cohort, aliquots of each of the individual samples were combined to make technical replicates, which were run as described above. Additionally, in each experiment, several buffer aliquots were run as blanks for artifact identification.

### Metabolomics data processing

Metabolite assignments to KEGG compounds were performed using MAVEN (Princeton, NJ, USA) on the basis of accurate intact mass, retention time and MS/MS^69,70^. Isotopologue distributions in tracing experiments with ^13^C^15^N-tryptophan were also analyzed with MAVEN and Compound Discoverer (Thermo Fisher, Bremen, Germany). Peak areas were exported for further statistical analysis with R (R Foundation for Statistical Computing, Vienna, Austria). Data sets from each of the two studies were log_2_ transformed. To perform differential expression analysis, the limma R package (v3.32.10) was used to fit a linear model to the data with age, sex, and cohort as covariates^35^. log_2_ fold change, p-value and adjusted p-value were calculated for the T21-D21 comparison using an unmoderated Student’s t-test and the false discovery rate (FDR) method for multiple testing correction^36^. Adjusted p-values are shown in the volcano plots and reported on the box plots. For displaying data per sample on the boxplots, we plotted the adjusted intensity values for each metabolite produced by the removeBatchEffects function in the limma R package using age, sex and cohort as covariates.

### RNAseq sample prep

Peripheral blood was collected in EDTA vacutainer tubes (BD, 366643) from ten individuals with T21 and nine D21 controls. Blood was centrifuged at 500 g for 15 min to separate plasma, buffy coat and red blood cells. White Blood Cells (WBCs) were isolated from the buffy coat fraction by RBC lysis and washed with our sorting buffer (see below) according to manufacturer’s instructions (BD, 555899). WBCs were cryopreserved in CryoStor CS10 Freezing Media (#210102) at ~10 million cells/mL. WBCs were awoken by quick thawing at 37 C and immediately spiked into 9 mL of sorting media and centrifuged at 500 g for 5 min and the media removed. Cells were lysed in 600 μL RLT plus (Qiagen) and Beta-mercaptoethanol (BME) lysis buffer (10 μL BME:1 mL RLT plus). RNA was extracted using the AllPrep DNA/RNA/Protein Mini Kit according to manufacturer’s instructions (Qiagen, 80004). RNA quality was determined by Agilent 2200 TapeStation and quantified by Qubit (Life Technologies). Samples with RIN of 6.8 or greater and a minimum of 1 μg were sent to Novogene for stranded library prep and 2×150 paired-end sequencing.

### RNAseq data analysis

Analysis of library complexity and high per-base sequence quality across all reads (i.e. q>30) was performed using FastQC (v0.11.5) software (Andrews 2010). Low quality bases (q<10) were trimmed from the 3’ end of reads and short reads (<30 nt after trimming) and adaptor sequences were removed using the fastqc-mcf (v: 1.05) tool from ea-utils. Common sources of sequence contamination such as mycoplasma, mitochondria, ribosomal RNA were identified and removed using FASTQ Screen (v0.9.1). Reads were aligned to GRCh37/hg19 using TopHat2 (v2.1.1, -- b2-sensitive --keep-fasta-order --no-coverage-search --max-multihits 10 --library-type fr-firststrand). High quality mapped reads (MAPQ>10) were filtered with SAMtools (v1.5). Reads were sorted with Picardtools (v2.9.4) (SortSAM) and duplicates marked (MarkDuplicates). QC of final reads was performed using RSeQC (v2.6.4). Gene level counts were obtained using HTSeq (v0.8.0,--stranded=reverse –minaqual=10 –type=exon –idattr=gene_id --mode= intersection-nonempty, GTF-ftp://igenome:G3nom3s4u@ussd-ftp.illumina.com/Homo_sapiens/UCSC/hg19/Homo_sapiens_UCSC_hg19.tar.gz). Differential expression was determined using DESeq2 (v1.18.1) and R (v3.4.3).

### Cell culture

Human fibroblasts were described previously^11^. For the metabolic flux experiment, cells were cultured in 5 mg/L ^13^C_11_ ^15^N_2_ L-tryptophan (Sigma-Aldrich, 574597) for 72 hours prior to stimulation with 10 ng/mL IFN-α (R&D Systems, 11101-2). Media was prepared by adding 120 μL of 10 mg/mL stock ^13^C_11_ ^15^N_2_ L-tryptophan in 1x PBS to 240 mL of RPMI (Corning, 10-040) in 10% Dialyzed FBS (Sigma, F0392) and 1x Anti-Anti (Gibco, 15240-062). Cells were treated with 10 ng/mL IFN-α or vehicle by adding 6 μL of 0.2 mg/mL IFN-α or 1x PBS to 120 mL of heavy labeled media. Cell lysates and media were collected at 0, 1, 6, and 24 hours following initial application of the heavy tryptophan and IFN-α. LC-MS metabolomics profiling was performed as described above, and tryptophan catabolites were identified by their ^13^C and/or ^15^N peaks.

### Generation of shRNA and CRISPR cell lines

To generate lentiviral particles 3.0e6 HEK293FT cells were transfected with 10 μg shIFNAR1 (TRCN0000059013, GCCAAGATTCAGGAAATTATT), IFNAR1 gRNA (Human GeCKO Library A 22695, AACAGGAGCGATGAGTCTGT), or IFNAR2 gRNA (Human GeCKO Library A 22697, GTGTATATCAGCCTCGTGTT) plasmid, and 10 μg of packaging viral mix (3:1 pD8.9:pCMV-VSVG) using PEI in a10 cm plate. 2 mL of freshly harvested lentiviral particles were then used to transduce 1.0e5 T21 cells in each well of a 6-well plate. The cells were expanded to 10 cm plates before starting puromycin selection (1 μg/mL).

### Fibroblast IFN stimulations and Western blots

Cells were plated at equal densities (~33,000/cm^2^) and allowed to attach overnight. The next day, IFN-α (R&D Systems, 11101-2) was added at 10 ng/mL in RPMI (Corning, 10-040) and cells were harvested by 0.25% trypsin (Gibco, 25200-056) at 0, 1, 6, and 24 hours. Trypsin was quenched by adding RPMI with 10% Dialyzed FBS (Sigma, F0392). Cells were pelleted at 200 g for 5 minutes. Cell pellets were washed with 1x PBS, then resuspended in RIPA buffer containing 1 μg/mL pepstatin, 2 μg/mL aprotonin, 20 μg/mL trypsin inhibitor, 10 nM leupeptin, 200 nM Na3VO4, 500 nM phenylmethylsulfonyl fluoride (PMSF), and 10 μM NaF. Suspensions were sonicated at six watts for 15 s two times and clarified by centrifugation at 21,000 g for 10 min at 4°C. Supernatants were quantified in a Pierce BCA Protein Assay and diluted in complete RIPA with 4x Laemmli sample buffer. Tris-glycine SDS-polyacrylamide gel electrophoresis was used to separate 20–40 μg protein lysate, which was transferred to a 0.45 μm polyvinylidene fluoride (PVDF) membrane. Membranes were blocked in 5% non-fat dried milk or 2.5% bovine serum albumin (BSA) in Tris-buffered saline containing 0.1% TWEEN (TBS-T) at room temperature for 30 min. Immunoblotting was done using primary antibodies against IDO1 (Cell Signaling Technology, 86630), and GAPDH (Santa Cruz Biotechnology, 365062) overnight in 5% non-fat dried milk or 2.5% BSA in TBS-T at 4°C while shaking. Membranes were washed 3x in TBS-T for 5–15 min before probing with a horseradish peroxidase (HRP) conjugated secondary antibody in 5% non-fat dried milk at room temperature for one hour. Membranes were again washed 3x in TBS-T for 5–15 min before applying enhanced chemiluminescence (ECL) solution. Chemiluminensence signal was captured using a GE (Pittsburgh, PA) ImageQuant LAS4000.

### Q-RT-PCR

Total RNA was harvested from cells using TRIzol (Life Technologies/Thermo Fisher Scientific) and reverse transcription was carried out using the Applied Biosystems High Capacity cDNA kit (Life Technologies/Thermo Fisher Scientific). Quantitative PCR was carried out with reference to a standard curve using SYBR Select Master Mix for CFX (Life Technologies/Thermo Fisher Scientific) on a Viia7 Real-Time PCR system (Life Technologies/Thermo Fisher Scientific), and normalized to 18S rRNA signals. Primers used for qRT-PCR were as follows (5’ to 3’): 18S-F, GCCGCTAGAGGTGAAATTCTTG; 18S-R, CTTTCGCTCTGGTCCGTCTT; IDO1-F, CCGTAAGGTCTTGCCAAGAAATA; IDO1-R, GTCAGGGGCTTATTAGGATCC; IFNAR1-F, GATTATCAAAAAACTGGGATGG; IFNAR1-R, CCAATCTGAGCTTTGCGAAATGG; IFNAR2-F, GGTTCTCATGGTGTATATCAGC; IFNAR2-R, GCAAGATTCATCTGTGTAATCAGG.

### Fibroblast flux data analysis

Prior to plotting, flux data was batch-adjusted and measurements from biological replicates were summarized on time-course plots as the mean value of the replicates +/− the standard error of the mean. P-values were calculated in R with a Student’s t-test and used the FDR method for multiple testing correction.

### Mesoscale discovery assay and correlation of cytokines to KYN levels

The Mesoscale Discovery Human Biomarker 54-Plex was combined with U-Plex reagents for IFN-α2a, IFN-β, and IL-29 (IFN-λ), to measure a total of 57 cytokines from 500 μL of plasma from each of the 67 individuals in Cohort 2 per manufacturer’s instructions. To visualize the effect of karyotype on each cytokine, the MSD data from the both the D21 and T21 samples were z-scored using the mean and standard deviation from the D21 group. Significantly differentially expressed cytokines were determined using the Kolmogorov-Smirnov test in R and an FDR-adjusted p-value threshold of 0.05. Prior to correlation analyses, both MSD data and metabolomics data were adjusted for age and sex using the linear model fitting procedure described above. Spearman’s rank correlations and p-values were calculated in R using the cor.test function. Adjusted p-values used FDR for multiple testing correction.

### Mouse strains and data analysis

Dp(10Prmt2-Pdxk)1Yey/J (Dp(10)1Yey/+), Dp(16Lipi-Zbtb21)1Yey/J (Dp(16)1Yey/+), and Dp(17Abcg1-Rrp1b)1(Yey)/J (Dp(17)1Yey/+)have been previously described^46^. Dp(16) mice were purchased from Jackson Laboratories or provided by Drs. Faycal Guedj and Diana Bianchi at the National Institutes of Health. Dp(10)1Yey/+, and Dp(17)1(Yey)/+ mice were provided by Drs. Katheleen Gardiner and Santos Franco, respectively. Animals were used between 6 and 30 weeks of age. All mice were maintained on a C57Bl/6 background and housed in specific-pathogen-free conditions. All experiments were approved by the Institutional Animal Care and Use Committee at the University of Colorado Anschutz Medical Campus. Prior to plotting and testing for statistical significance, mouse metabolomics data was adjusted for batch and sex using the linear modeling procedure described above. P-values were calculated in R with a Student’s t-test and used FDR for multiple testing correction.

## Supporting information

Supplementary Figures and Data Descriptions

## Data and code availability

All data and R code developed for the full set of statistical and machine learning analyses is available at: https://github.com/CostelloLab/Trisomy21_KYN_metabolomics. To ensure reproducibility, this code can be run to generate all figures for this manuscript.

## Acknowledgements

This work was supported primarily by the Linda Crnic Institute for Down Syndrome, the Global Down Syndrome Foundation, the Anna and John J. Sie Foundation, the Boettcher Foundation, and the University of Colorado School of Medicine. We thank the University of Colorado Cancer Center Human Immune Monitoring Shared Resource for assistance with Mesoscale Discovery Assay studies. We thank the individuals with Down syndrome that donated the biological samples that enabled these studies.

## ADDITIONAL INFORMATION

### Author Contributions

RKP, KDS, JME, and JCC, conception and design, acquisition of data, analysis and interpretation of data, drafting or revising the article; AD, conception and design, acquisition of data, analysis and interpretation of data; RCH, MPL, KPS, KAW, RM, KDT, HCL, ALR, REG, RBW, DEK, MJ, acquisition of data, analysis, and interpretation of data.

