## Supplementary Figures and Data Descriptions for "Trisomy 21 activates the kynurenine pathway via increased dosage of interferon receptors"

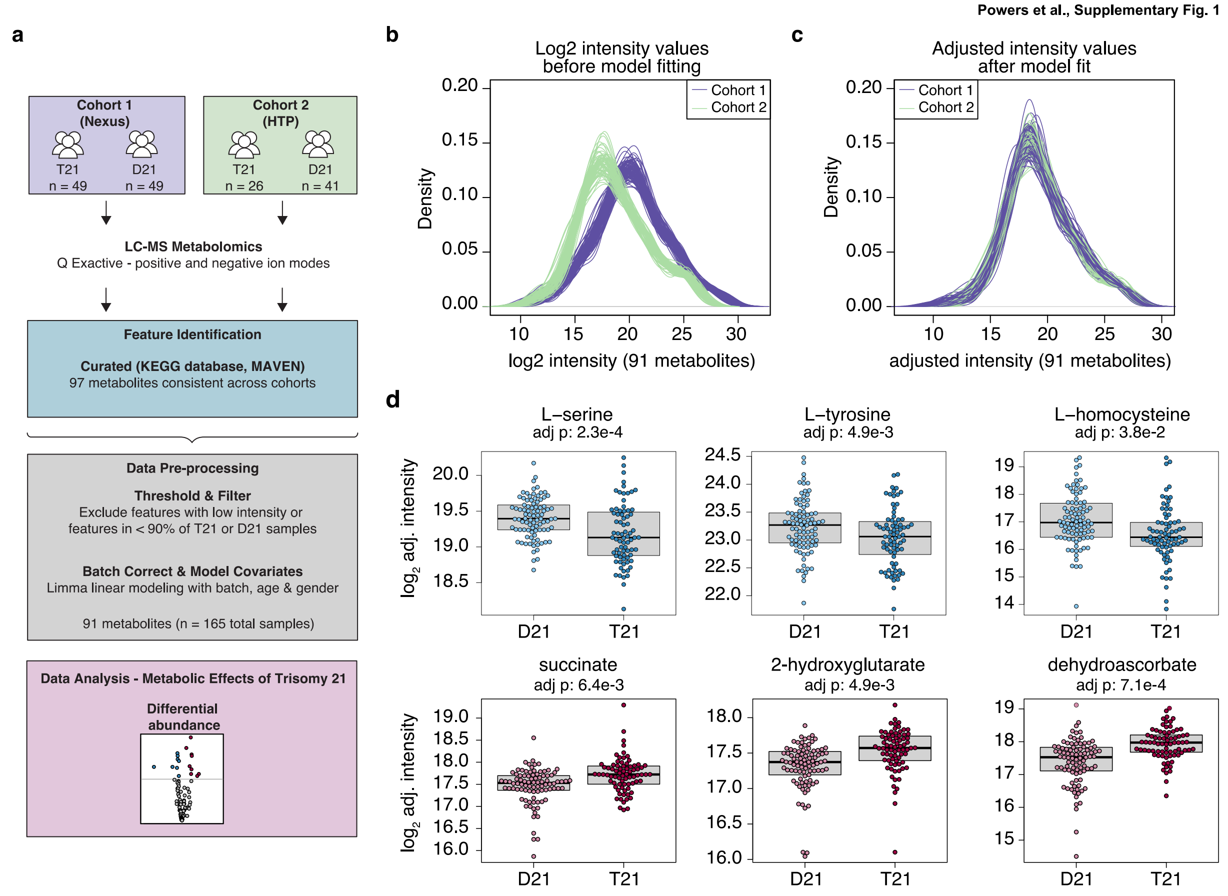


### Supplementary Figure 1 - Statistical workflow adjusts for the effects of age and sex in circulating plasma metabolites.

**a)** Overview of the analysis pipeline used to identify, annotate, and analyze plasma metabolites measured in individuals with Down syndrome (T21 group) and controls without Down syndrome (D21) from the Translational Nexus Biobank (Nexus, Cohort 1) and the Human Trisome Project (HTP, Cohort 2). Putative metabolites were annotated using MAVEN. Across the two cohorts, 97 metabolites were detected consistently. Data preprocessing included filtering out metabolites with low intensity values or that were detected in less than 90% of the T21 and/or D21 samples, leaving 91 metabolites. Batch correction and linear model fitting was performed before the combined data set was used for analysis. **b)** Density plots showing the distribution of $\log_2$ intensity values for all 91 metabolites in Cohort 1 (purple) and Cohort 2 (green) prior to any adjustment. **c)** Density plots showing the distribution of adjusted $\log_2$ intensity values for all 91 metabolites, colored as in **b**, after a linear model was used to adjust for age, sex, and cohort covariates. **d)** Boxplots showing $\log_2$ adjusted intensities for significantly differentially abundant metabolites. p-values were calculated using the linear model from **a** and the FDR method for multiple testing correction.

**a**

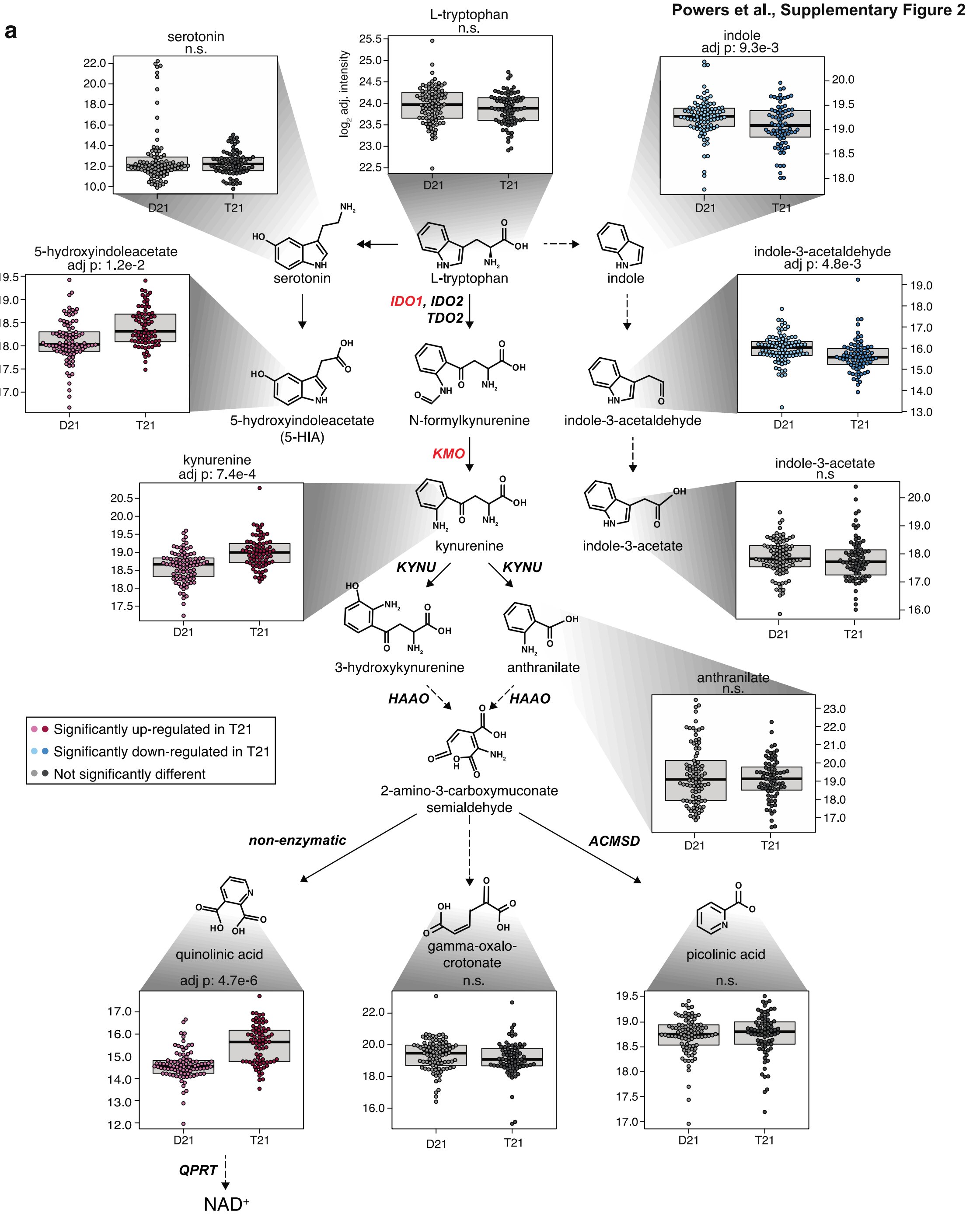


**b**

RNAseq - white blood cells

| Gene | $log_2$foldchange | adj p |
|---|---|---|
| *IDO1* | 1.24 | 8.04e-5 |
| *IDO2* | 1.27 | 3.85e-1 |
| *TDO2* | - | - |
| *KMO* | -0.58 | 7.71e-4 |
| *KYNU* | 0.00 | 9.97e-1 |
| *HAAO* | -0.41 | 3.32e-1 |
| *QPRT* | 0.24 | 5.30e-1 |
| *ACMSD* | - | - |
| *KYAT1* | - | - |
| *AADAT* | - | - |
| *KYAT3* | - | - |

**c**

RNAseq - white blood cells

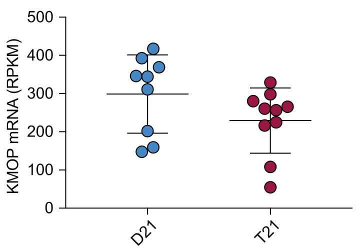

**Supplementary Figure 2 – Trisomy 21 induces the kynurenine pathway. a)** Schematic of tryptophan metabolism and the kynurenine pathway. Metabolite levels are plotted as model residuals (i.e. remaining component of the model fit after adjusting for age, sex, and cohort). p-values were calculated using the linear model from **Figure 1a** with FDR for multiple testing correction. Multi-headed arrows are used to represent multiple steps in the pathway that are not shown. Arrows with dashed lines represent reactions occurring in tryptophanase-expressing gastrointestinal microbiota. **b)** Table displaying fold change and significance level for each enzyme in the kynurenine pathway. Data are derived from mRNA-seq of white blood cells from individuals with trisomy 21 and controls. Absent values indicate no detectable mRNA expression. **c)** Scatter plot showing RNA expression of *KMO* in white blood cells from controls and individuals with T21. Statistical significance was calculated using DESeq2. mRNA expression values are displayed in reads per kilobase per million (RPKM)**.**

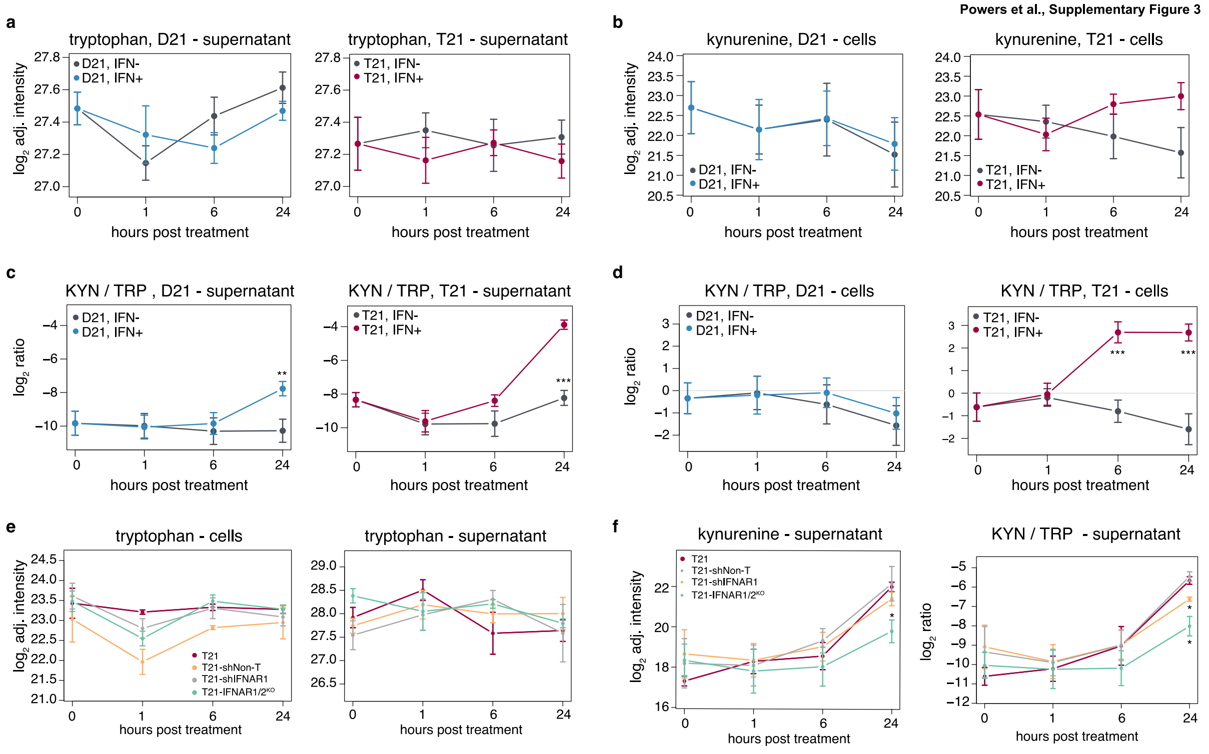


**Supplementary Figure 3 – Trisomy 21 sensitizes cells to super-induction of the kynurenine pathway by IFN-α.**

Metabolic flux experiment using heavy-labeled ($^{13}C^{15}N$) tryptophan in fibroblast cell lines (n = 3 T21, n = 3 D21). **a)** Levels of heavy-labeled tryptophan in D21 (left) and T21 (right) supernatant, with or without IFN-α treatment. **b)** Levels of heavy-labeled kynurenine in D21 (left) and T21 (right) fibroblast cell lysates, with or without IFN-α treatment. **c)** Ratio of heavy-labeled kynurenine to heavy-labeled tryptophan levels in D21 (left) and T21 (right) fibroblast supernatants, with or without IFN-α treatment. **d)** Ratio of heavy-labeled kynurenine to heavy-labeled tryptophan levels in D21 (left) and T21 (right) fibroblast cell lysates, with or without IFN-α treatment. At each timepoint, the p-value between the Ctrl and IFN+ samples was calculated using a two-tailed student's t-test (* $p < 0.05$, ** $p < 0.01$, *** $p < 0.001$). **e.** Levels of heavy-labeled tryptophan in the indicated cells (left) and supernatants (right) during a 24 hour time course of IFN-α treatment. **f.** Levels of heavy-labeled kynurenine in the supernatants of the indicated cell line (left) and ratio of heavy-labeled kynurenine to heavy-labeled tryptophan levels in the supernatants of the indicated cell line during a 24 hour time course of IFN-α treatment. At each timepoint, the p-value between the parental T21 cell line and each T21 cell line with modified *IFNAR* levels was calculated using a two-tailed student's t-test (* $p < 0.05$, ** $p < 0.01$, *** $p < 0.001$).

**Supplementary Data.**

**Supplementary Data 1. Cohort Details.** Demographic data for each of the three cohorts studied in this manuscript. **Column: a)** Sample ID, **b)** Sex, **c)** Age, **d)** Karyotype.

**Supplementary Data 2. Metabolomics Data.** Differential abundance analysis of 91 metabolites in both cohorts using Limma linear model fitting with an unmodified t-test. **Column: a)** Compound ID, **b)** Compound Name, **c-e** and **g-i**, $Log_2$ Fold Change, p-value, and Adjusted p-value for Batch Covariate, and Batch, Age and Sex Covariates, respectively.

**Supplementary Data 3. White Blood Cell RNA-seq.** Differential gene expression analysis of white blood cell RNA-seq data from individuals with trisomy 21 and controls using DESeq2. **Column: a)** GeneID, **b)** Chromosome, **c)** baseMean, **d)** baseMeanD21, **e)** baseMeanT21, **f)** Fold Change, **g)** $Log_2$ Fold Change, **h)** Adjusted p-value.

**Supplementary Data 4. Mesoscale Discovery Assay.** Differential abundance analysis of 55 cytokines from Cohort 2 using Kolmogorov-Smirnov test with FDR correction**. Column: a)** Cytokine, **b)** Kolmogorov-Smirnov D value, **c)** $Log_2$ Fold Change, **d)** p-value, **e)** Adjusted p-value.

**Supplementary Data 5. Spearman's Rank Correlations.** Correlation analysis of kynurenine and 55 cytokines using Spearman's rank correlation. **Column: a)** Cytokine, **b-d), e-g), h-j),** Spearman's rho ($\rho$), p-value and Adjusted p-value for all individuals, only T21 individuals, and only control individuals, respectively.

**Supplementary Data 6. $^{13}C^{15}N$-tryptophan Metabolic Flux Data.** Differential abundance analysis of $^{13}C^{15}N$-tryptophan metabolites using Limma linear model fitting with an unmodified t-test. **Tab a)** Metabolic flux analysis of six fibroblasts cell lines (n = 3 T21, n = 3 D21) upon IFN-α treatment. **Column: a)** Sample Type, **b)** Karyotype, **c)** Timepoint, **d)** Compound, **e)** Comparison, **f)** Log2 Fold Change, **g)** p-value. **Tab b)** Metabolic flux analysis of IFNAR knockdown or knockout fibroblast cell lines upon IFN-α treatment. **Column: a)** Sample Type, **b)** Cell Line, **c)** Timepoint, **d)** Treatment, **e)** Compound, **f)** Comparison, **g)** Log2 Fold Chang, **h)** p-value.

**Supplementary Data 7. Mouse Metabolomics Data.** LC-MS metabolomics of KYN levels in WT, Dp16, Dp10, Dp17 mouse strains. **Column: a)** Mouse Strain, **b)** Batch, **c)** Sex, **d)** $Log_2$ Unadjusted KYN Intensity and **e)** Batch- and Sex-adjusted $Log_2$ KYN Intensity.
